# The critical role of ASD-related gene *CNTNAP3* in regulating synaptic development and social behavior in mice

**DOI:** 10.1101/260083

**Authors:** Da-li Tong, Rui-guo Chen, Yu-lan Lu, Wei-ke Li, Yue-fang Zhang, Jun-kai Lin, Ling-jie He, Ting Dang, Shi-fang Shan, Xiao-Hong Xu, Yi Zhang, Chen Zhang, Ya-Song Du, Wen-Hao Zhou, Xiaoqun Wang, Zilong Qiu

## Abstract

Accumulated genetic evidences indicate that the contactin associated protein-like (CNTNAP) family is implicated in autism spectrum disorders (ASD). In this study, we identified genetic mutations in the *CNTNAP3* gene from Chinese Han ASD cohorts and Simons Simplex Collections. We found that CNTNAP3 interacted with synaptic adhesion proteins Neuroligin1 and Neuroligin2, as well as scaffolding proteins PSD95 and Gephyrin. Significantly, we found that CNTNAP3 played an opposite role in controlling the development of excitatory and inhibitory synapses *in vitro* and *in vivo,* in which ASD mutants exhibited *loss-of-function* effects. In this study, we showed that *Cntnap3-null* mice exhibited deficits in social interaction, spatial learning and prominent repetitive behaviors. These evidences elucidate the pivotal role of CNTNAP3 in synapse development and social behaviors, providing the mechanistic insights for ASD.

## Introduction

Autism spectrum disorder (ASD) is a prevalent neurodevelopmental disorder with early onset in the childhood, characterized by deficits in social behaviors and prominent repetitive behaviors. Numerous genes have been discovered to associate with ASD by human genetic studies(de la Torre-Ubieta et al., 2016; Huguet et al., 2013; Willsey and State, 2015). Notably, it has been reported that mutations in genes encoding synaptic adhesion molecules, including neuroligin (NLGN) and neurexin (NRXN) family members, are closed related to ASD, which are often found in ASD patients, suggesting that the synaptic dysfunction significantly contribute to ASD(de la Torre-Ubieta et al., 2016; Huguet et al., 2013; Willsey and State, 2015).

As a member of NRXN superfamily, the contactin associated protein-like (CNTNAP) family (also known as the CASPR protein family) has been identified to be associated with ASD, especially CNTNAP2 and CNTNAP4. The CNTNAP family contains 5 members from CNTNAP1 to CNTNAP5, featured by multiple repeats of epidermal growth factor (EGF) domains and laminin G (LamG) domains in the extracellular domains, as well as the intracellular PDZ-binding domain (Bellen et al., 1998; Peles et al., 1997; Spiegel et al., 2002; Traut et al., 2006). CNTNAP1 and CNTNAP2 are involved in the formation of myelin and trafficking potassium channels of the cell membrane (Poliak et al., 2003; Rios et al., 2000; Traka et al., 2003). Genetic studies suggested that the CNTNAP2 gene had strong connections with ASD (Alarcon et al., 2008; Arking et al., 2008; Bakkaloglu et al., 2008). The CNTNAP2 protein plays a critical role in neural development and synaptic transmission. Furthermore, *Cntnap2* null mice exhibited various autistic-like behaviors (Anderson et al., 2012; Penagarikano et al., 2011). CNTNAP4 is fateful for the formation of axo-axonic synapse by interacting with Contactin5 in the peripheral nervous system (Ashrafi et al., 2014). Whereas in the central nervous system, CNTNAP4 plays a vital part in regulating synaptic transmission of GABAergic neurons. Moreover, the *Cntnap4* knockout mice also showed heavily stereotypic behaviors like self-grooming (Karayannis et al., 2014).

The whole-genome sequencing of 476 autism families from the Simons Simplex Collection detected a stop-gain *de novo* mutation in CNTNAP3 (R1219X) in an ASD proband (Turner et al., 2017). Notably, CNTNAP3 was also found to express differentially in the blood of individuals with autism (Kong et al., 2012). Furthermore, a deletion of 9p12 which contains *CNTNAP3* was found in the mental retardation (MR) patient (Mosrati et al., 2012). Taken together, CNTNAP3 may be a candidate gene for autism spectrum disorders.

In this study, we identified genetic mutations in the *CNTNAP3* gene through whole-exome sequencing in Chinese Han ASD cohorts and Simons Simplex Collections. To address molecular mechanisms underlying CNTNAP3 regulating synaptic development and social behaviors, we performed extensive molecular, physiological and genetic experiments. We found that CNTNAP3 interacted with critical synaptic adhesion molecules such as Neuroligin1 and Neurogligin2 proteins, as well as postsynaptic scaffolding protein PSD95 and Gephyrin. More importantly, our data demonstrated that CNTNAP3 played an opposite role in controlling development of excitatory and inhibitory synapses *in vitro* and *in vivo,* in which ASD mutants exhibited *loss-of-function* effects. Furthermore, *Cntnap3^-/-^* mice exhibited deficits in social behavior, cognitive tasks and prominent repetitive behaviors, confirming the role of CNTNAP3 in ASD.

## Results

### Identification of *CNTNAP3* mutations in ASD patients

In the whole-exome sequencing study of 120 ASD patients collected from Shanghai Mental Health Hospital (Wen et al., 2017), we found two probands (ASD22, ASD652) carrying the same inherited mutation (causing amino acid alternation P614A) in the *CNTNAP3* gene, validated by Sanger sequencing (Fig. 1a, Supplementary Fig. 1a, d, e). We identified another proband featured “Human phenotype ontology (HPO): autism” in the database of brain disorders patients in Fudan Children’s Hospital of Shanghai, in which an inherited mutation (R786C) transmitted from the father in the *CNTNAP3* gene were found, suggesting the connection between *CNTNAP3* and ASD (Fig. 1a, Supplementary Fig. 1b, f).

**Figure 1.**
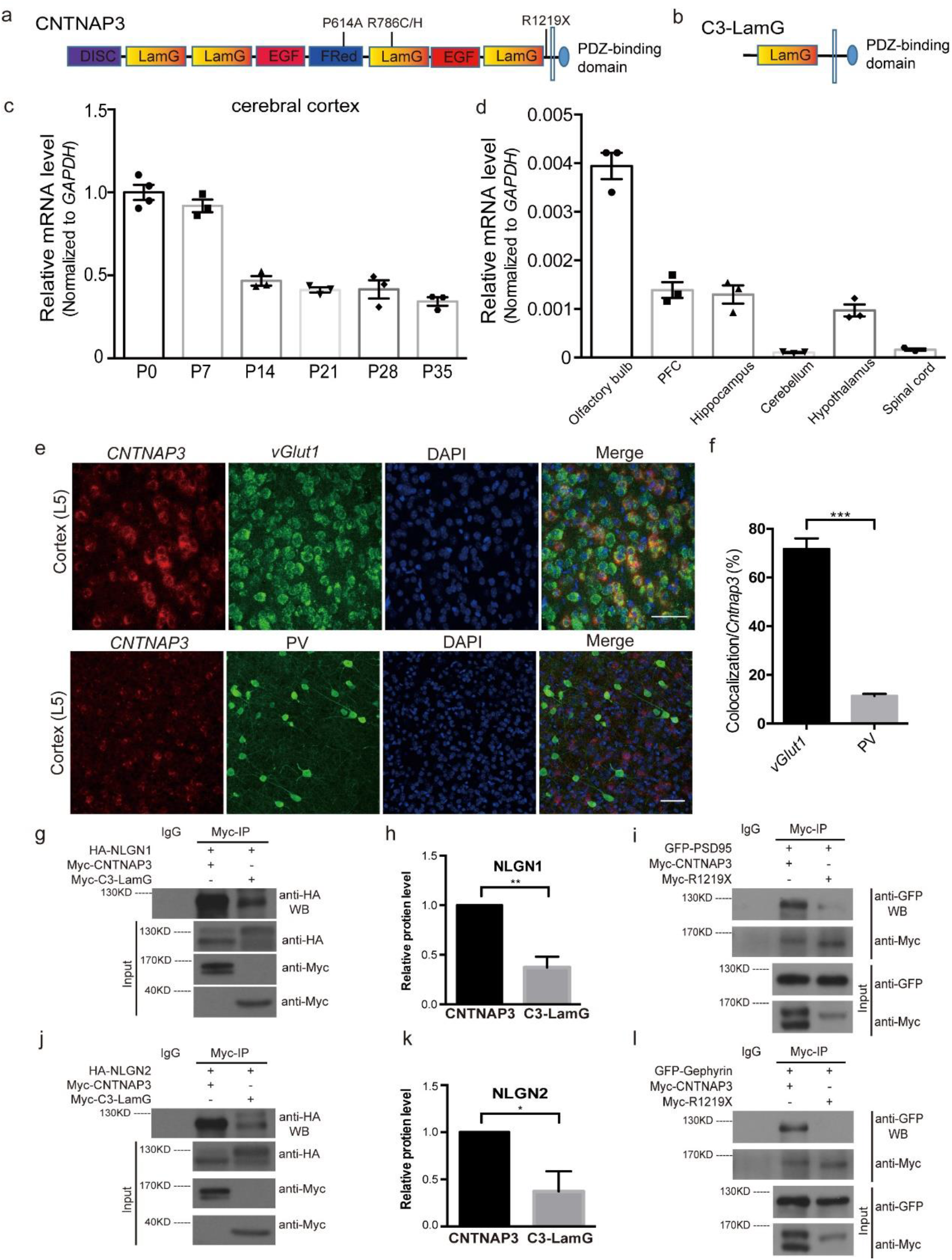
ASD-related CNTNAP3 mutations and the expression pattern of Cntnap3 in mouse brain. (a) The schematic structure and locations of P614A, R786C/H and R1219X in the CNTNAP3 protein. (b) The schematic structure of CNTNAP3-LamG (C3-LamG for short). (c) Expression levels of *Cntnap3* in cerebral cortex of mouse in different ages (C57BL/6). (d) Expression levels of *Cntnap3* in different regions of the CNS of mouse (C57BL/6, Postnatal 28 days: P28). (e) Double labeling with immunostaining and *in situ* hybridization on P14 wild-type cortex with co-localization of *Cntnap3 (in situ,* red) and *vGlut1 (in situ,* green) or PV (immunostaining, green). (f) Mean percentage of co-localization of *Cntnap3* and *vGlut1* or PV (co-localization / *Cntnap3* positive). (g) Co-immunoprecipitation of myc-CNTNAP3, myc-C3-LamG and HA-NLGN1 in 293T cells. Immunoprecipitated (IP) with anti-myc antibody, immunoblotted (IB) with anti-myc and anti-HA antibodies. (h) Quantitative analysis of HA-NLGN1 protein level interacts with myc-CNTNAP3 or myc-LamG. (i) Co-immunoprecipitation of myc-CNTNAP3, myc-R1219X and GFP-PSD95 in 293T cells. IP with anti-myc antibody, IB with anti-myc and anti-GFP antibodies. (j) Co-immunoprecipitation of myc-CNTNAP3, myc-C3-LamG and HA-NLGN2 in 293T cells. IP with anti-myc antibody, IB with anti-myc and anti-HA antibodies. (k) Quantitative analysis of HA-NLGN2 protein level interacts with myc-CNTNAP3 or myc-LamG. (l) Co-immunoprecipitation of myc-CNTNAP3, myc-R1219X and GFP-Gephyrin in 293T cells. IP with anti-myc antibody, IB with anti-myc and anti-GFP antibodies. (c, d, mouse number n = 3, Scale bar: 50μm.) Statistical significance was evaluated by Student’s t test: * p < 0.05, ** p < 0.01; error bars, ± SEM.

To determine whether *CNTNAP3* mutations exist in ASD populations from different geographic regions, we next searched the Simons simplex collections (SSC) and found 2 ASD probands carrying R786H mutation in the *CNTNAP3* gene, among 2600 ASD trios, both of which are inherited from unaffected mothers (Supplementary Fig. 1c). In conclusion, there are 5 mutations of CNTNAP3 (G410S, P614A, R786C/H, R1219X) reported to be found in ASD patients yet, which caused 4 amino acid changes and one stop-gain *de novo* mutation (Vaags et al., 2012). We further examined the mutation rates of *CNTNAP3* mutations (G410S, P614A, R786C, R786H and R1219X) in the gnomAD database (http://gnomad.broadinstitute.org). P614A, R786C and R1219X exhibited extremely low occurrence (less than 0.01%) in total 245686 populations of the gnomAD database, indicating that they are rare variants. The R786H mutation occurs 0.012% in gnomAD database, which is still much lower than it in SSC (2/2600, 0.077%), suggesting the possible enrichment of R786H mutation in ASD cohorts. However, G410S exhibited high occurrence (3.7%) in total 276826 populations of the gnomAD database, indicating that G410S may be just a single-nucleotide polymorphism. The R786 locates in the LamG domains, P614 is within fibrinogen-related (FRed) domain, and R1219 locates in the link region before trans member domain (Fig. 1a).

### CNTNAP3 interacts with NLGNs and NRXNs

To investigate how CNTNAP3 participate in brain development, we first examined the expression profile of CNTNAP3 in developmental stages and various regions in the mouse brain via quantitative PCR (Fig. 1c, d), due to lack of the suitable antibody against the mouse CNTNAP protein. We found that CNTNAP3 was highly expressed in cortex and hippocampus of the mouse postnatal brain (Fig. 1c, Supplementary Fig. 2a) and had relative low expression levels in cerebellum and spinal cord (Fig. 1d). We further performed *in situ* hybridization using probes against the mouse *Cntnap3* gene together with immunostaining with various cell markers in cortex and hippocampus to characterize the expression pattern of CNTNAP3 in different neuronal cell types (Fig. 1e, Supplementary Fig. 2c). Interestingly, we found that the majority (~70%) of CNTNAP3-expressing neurons were co-localized with vesicular glutamate transporter 1 (vGlut1) in cerebral cortex, whereas around 10% CNTNAP3-expressing neurons are parvalbumin (PV)-positive, indicating that CNTNAP3 is mainly expressed in glutamatergic neurons in cortex (Fig. 1f).

CNTNAP3 has repeated LamG domains and EGF domains, as well as an intracellular PDZ-binding domain, which is structurally similar with NRXN superfamily proteins (Fig. 1a), hence, we would like to determine whether CNTNAP3 interact with synaptic adhesion molecules, such as Neuroligin1 (NLGN1), Neuroligin 2 (NLGN2) and synaptic scaffolding proteins. We constructed a plasmid expressing hemagglutinin (HA)-tagged full-length human CNTNAP3. To address which domain would be responsible for the interaction, we also made the plasmid expressing only one extracellular LamG domain and the intracellular PDZ-binding domain of CNTNAP3 protein (C3-LamG) (Fig. 1b).

By the co-immunoprecipitation experiment, we found the CNTNAP3 interacted with both NLGN1 and NLGN2 (Fig, 1g, j). Deletion of extracellular LamG domains (C3-LamG) significantly weaken, but not block the interaction, indicating that only one LamG domain is sufficient for the interaction between CNTNAP3 and NLGN1 or NLGN2 (Fig. 1h, k). CNTNAP3 also interacted with synaptic scaffolding proteins, including PSD-95 and Gephyrin but not SHANK3 (Fig. 1i, l, Supplementary Fig. 3a). NLGN1 and PSD-95 primarily located in the post-synaptic compartment of excitatory synapse, whereas NLGN2 and Gephyrin localized in the inhibitory synapses, these evidences suggest that CNTNAP3 may exist in both excitatory and inhibitory synapses.

Among the 5 mutations, R1219X has almost all of the extracellular region but not trans member region or PDZ-binding domain. We presumed that R1219X cloud not interact with PSD-95 or Gephyrin because of the absent of PDZ-binding domain (Fig. 1i, l), which suggested R1219X may be a *loss of function* mutation with the disability of recruiting scaffolding proteins.

### CNTNAP3 regulates excitatory and inhibitory synapse development differentially

To assess the function of CNTNAP3 in synaptic development, we constructed a short hairpin RNA (shRNA) against the mouse *Cntnap3* gene and confirmed the efficiency of down-regulating the endogenous expression level of *Cntnap3* (Supplementary Fig. 3b). We next investigated whether the knockdown of *Cntnap3* could affect neuronal morphology and synaptic development in cultured mouse primary neurons. We transfected GFP expressing plasmids, together with control vector, *Cntnap3* shRNA, human WT *CNTNAP3* cDNA as well as human *CNTNAP3* cDNA carrying ASD-related mutations, respectively. Since mouse *Cntnap3* shRNA does not target human *CNTNAP3,* human WT *CNTNAP3* cDNA could serve as a rescue construct for shRNA knockdown. We found that knockdown of CNTNAP3 expression led to significantly decrease of axonal and dendritic length, as well as dendritic branch number (Fig. 2a-c, Supplementary Fig. 4). It is worth noting that although human WT *CNTNAP3* cDNA was able to fully restore axonal and dendritic length to normal level after shRNA knockdown, three ASD-related mutations (R1219X, P614A and R786C) were not able to rescue defects caused by *Cntnap3* shRNA, exhibiting *loss-of-function* effects (Fig. 2b, c, Supplementary Fig. 4b, d).

**Figure 2.**
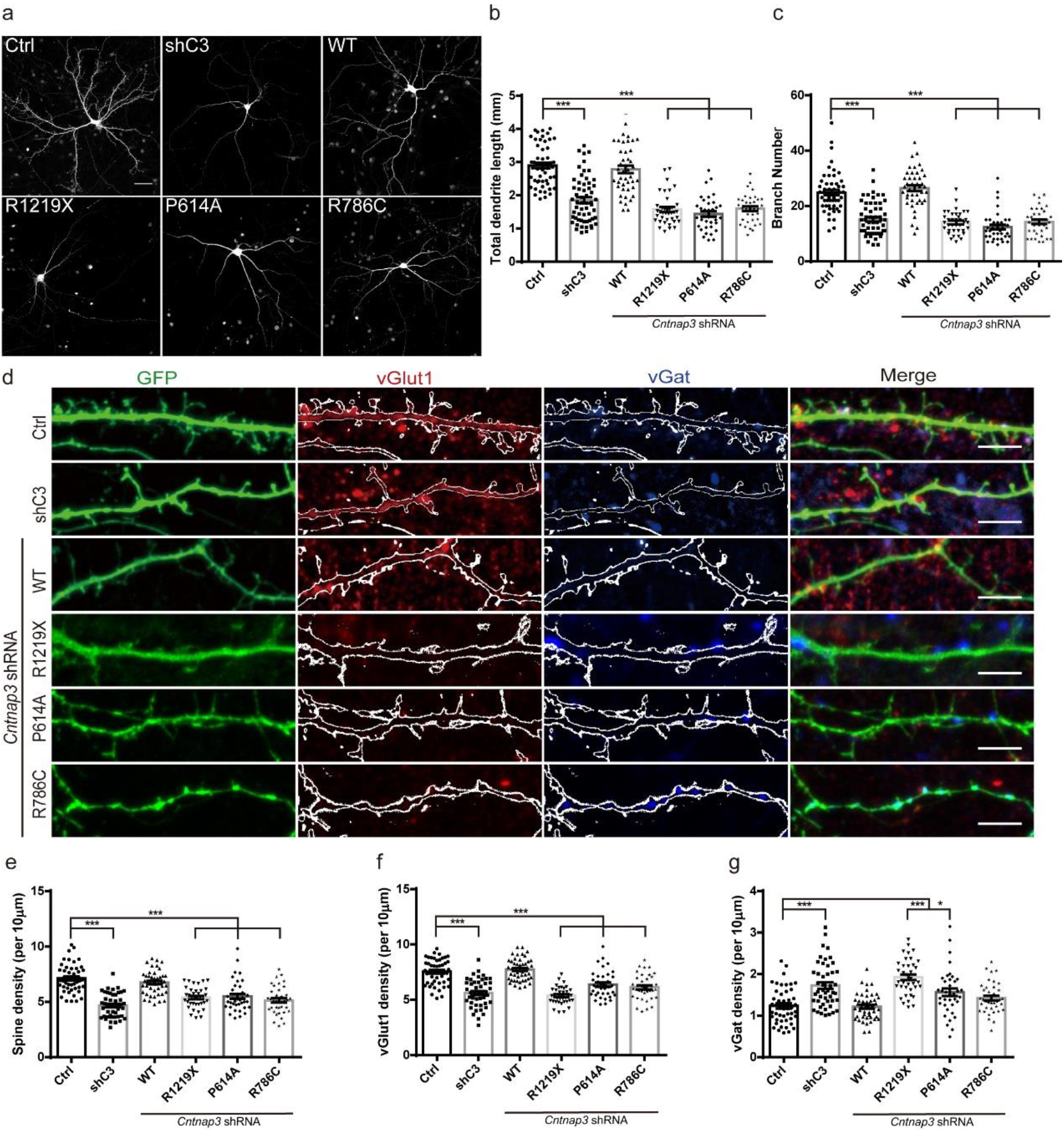
CNTNAP3 regulated neurite and synapse development. (a) Immunostaining of GFP (green) in cultured E14.5 cortex neurons. (E14.5+DIV 14, Scale bar: 50μm) (b) Dendritic length analysis of (a). (Counted neuron numbers Ctrl: n=54, shC3: n=58, WT: n=43, R1219X: n=37, P614A: n=36, R786C: n=36.) (c) Branch numbers analysis in (a). (d) Immunostaining of GFP (green), vGlut1 (red) and vGat (blue) in cultured E14.5 cortex neurons. (E14.5+DIV 14, counted neuron numbers Ctrl: n=47, shC3: n=47, WT: n=47, R1219X: n=37, P614A: n=37 R786C: n=37. Scale bar: 5μm) (e) Quantitative analysis of spine density in (d). (f) Quantitative analysis of vGlut1 density in (d). (g) Quantitative analysis of vGat density in (f). Statistical significance was evaluated by one-way ANOVA: *p < 0.05, ** *p* < 0.01, *** *p* < 0.001. Error bars, ±SEM.

Next, we would like to investigate whether CNTNAP3 contributes to synaptic development. After transfections of GFP expressing plasmids, along with vector control, or *Cntnap3* shRNA, as well as *Cntnap3* shRNA with human WT *CNTNAP3* or ASD-related mutations, respectively in mouse primary cortical neurons *in vitro* for 14 days, we then measured amounts of excitatory synapses by anti-vesicular glutamate transporter 1 (vGlut1) immunostaining and numbers of inhibitory synapses by anti-vesicular GABA transporter (vGat) immunostaining onto the GFP-expressing neurons transfected with either control vectors or various manipulations (Fig. 2d). We found that knockdown of *Cntnap3* led to decrease of excitatory synapse formation which were fully rescued by WT CNTNAP3, but not the R1219X, P614A and R786C mutants (Fig. 2d, f). Interestingly, inhibitory synapse numbers measured by anti-vGat staining were increased after knockdown of CNTNAP3, which could be fully rescued to normal level by WT and R786C, but not R1219X nor P614A mutant, suggesting that CNTNAP3 may play a negative role in repressing formation of inhibitory synapse and R1219X, P614A may exhibit *loss-of-function* effects in this regard (Fig. 2d, g). Furthermore, numbers of spine were also markedly decreased in the CNTNAP3 knockdown group, which were restored to the normal level when co-expressed with WT CNTNAP3, but not R1219X, P614A nor R786C mutants (Fig. 2e). These evidences indicate that CNTNAP3 may promote excitatory glutamatergic synapse formation, whereas inhibit GABAergic synapse formation. Additionally, ASD-related mutations may exhibit *loss-of-function* effects regarding to excitatory and inhibitory synapses formation specifically.

### CNTNAP3 regulates development of excitatory synapse and PV-positive inhibitory neurons *in vivo*

To address the role of CNTNAP3 *in vivo,* we constructed *Cntnap3^-/-^* mouse by CRISPR/Cas9 technology and conditional knockout of *Cntnap3* mouse by flanking exon 3 with LoxP cassettes with homology recombination strategy (Supplementary Fig. 3c, d). We first examined whether development of excitatory synapses can be altered in *Cntnap3^-/-^* mice by measuring the morphology of dendritic spine of pyramidal neurons in hippocampal CA1 and S1 (primary somatosensory cortex) region. Golgi staining showed that spine density of hippocampal CA1 and S1 neurons significantly decreased in *Cntnap3^-/-^* mice in comparison with WT mice, suggesting that CNTNAP3 plays a critical role in excitatory synapse development *in vivo* (Fig, 3a, b). Next, we crossed *Nex-Cre* mice with *Cntnap3* ^flox/flox^ mice to specifically delete *Cntnap3* in excitatory neurons(Kashani et al., 2006). In *Nex-Cre: Cntnap3*^flox/flox^ mice, spine density also dramatically decreased in hippocampal CA1 neurons comparing to WT mice, indicating that CNTNAP3 likely regulates development of excitatory synapse through cell-autonomous manner (Fig. 3c, d).

**Figure 3.**
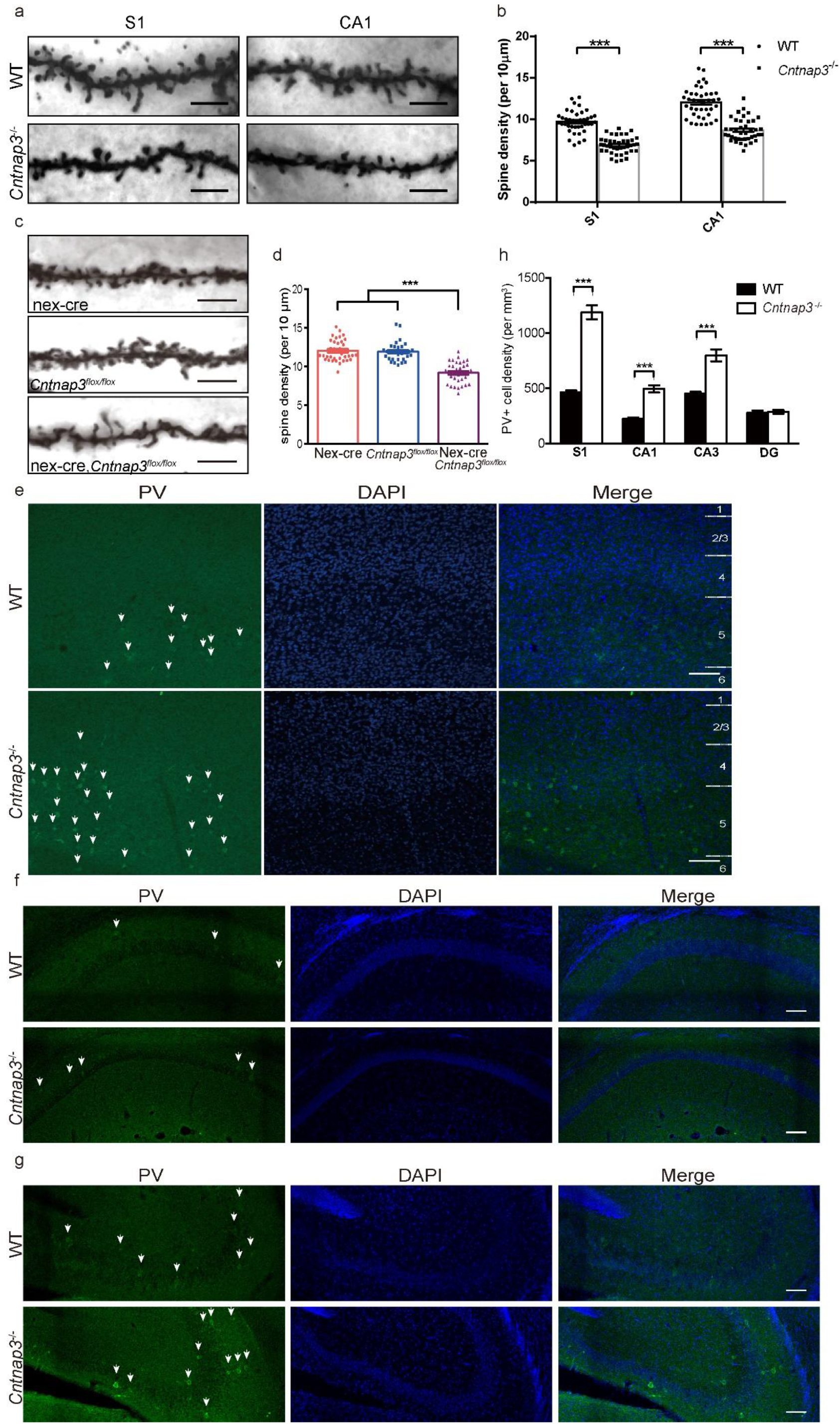
*Cntnap3^-/-^* mice showed decreased spine density and increased PV-positive neurons in cortex and hippocampus. (a) Golgi staining in adult *Cntnap3^-/-^* mice in S1 (primary somatosensory cortex) and CA1 region. (apical dendrite, 1 months of age, mice number n=3 for either WT or KO, scale bar: 5μm) (b) Quantitative analysis of spine density in *Cntnap3^-/-^* mice. (neuron numbers, WT: n=40, *Cntnap3^-/-^*: n=40 for both S1 and CA1 region) (c) Golgi staining in adult *Nex-Cre:Cntnap3*^flox/flox^ mice in CA1 region. (apical dendrite, from 2-4 months of age, mice number n=3 for each genotype, scale bar: 5μm) (d) Quantitative analysis of spine density in *Nex-Cre:Cntnap3*^flox/flox^ mice. (neuron numbers *Nex-Cre:* n=36, *Cntnap3*^flox/flox^: n=30, *Nex-Cre:Cntnap3*^flox/flox^: n=34, mice number n=3 for each genotype) (e) Immunochemistry of PV (green) in S1 region of *Cntnap3* KO mice at postnatal 14 days of age (P14). (Scale bar: 100μm) (f) Immunochemistry of PV (green) and in hippocampus CA1 region of P14 *Cntnap3^-/-^* mice. (Scale bar: 100μm) (g) Immunochemistry of PV (green) and in hippocampus CA3 region of P14 *Cntnap3^-/-^* mice. (Scale bar: 100μm) (h)Quantitative analysis of density of PV positive neurons in *Cntnap3^-/-^* mice. (mice number, n=3 for WT and KO) Statistical significance was evaluated by Student’s *t* test (*Cntnap3^-/-^* vs. WT) or one-way ANOVA (*Nex-Cre:Cntnap3^flox/flox^* vs. other groups): ***p < 0.001. Error bars, ± SEM.

Strikingly, we found that parvalbumin-positive GABAergic neurons markedly increased in cortical regions and hippocampus of *Cntnap3^-/-^* mice in comparison with WT mice (Fig. 3e-h, Supplementary Fig. 5a, b). But somatostatin (SST)-positive GABAergic neurons remained unaffected in *Cntnap3^-/-^* mice (Supplementary Fig. 5c and 6). These data suggest that CNTNAP3 may specifically hamper development of PV-positive subtype of GABAergic neurons.

### CNTNAP3 regulates excitatory and inhibitory synaptic transmission *in vivo*

To determine the role of CNTNAP3 in synaptic transmission *in vivo*, the whole-cell patch clamp recording was performed on the hippocampal CA1 neurons and layer 5 neurons of neocortex in acute slice preparations of *Cntnap3^-/-^* mice and WT mice as control (Fig. 4a). We firstly measured passive electrical properties of these cells, including resting membrane potential (RMP), input resistance (Rin) and membrane capacity (Cm) (Fig. 4b-d). Our results showed that intrinsic properties of neurons in layer 5 of *Cntnap3^-/-^* mice were equivalent to their counterparts in control mice (The upper panel of Fig. 4b-d). However, hippocampal CA1 pyramidal neurons of *Cntnap3^-/-^* mice appeared more depolarized compared to the cells in WT mice (The lower panel of Fig. 4b). Moreover, the Rin was elevated and the Cm decreased in *Cntnap3^-/-^* mice comparing to WT mice (The lower panel of Fig. 4c, d). These results suggest that membrane properties and ion channel distributions may be altered in hippocampal CA1 neurons in *Cntnap3^-/-^,* in the absence of *Cntnap3.*

**Figure 4.**
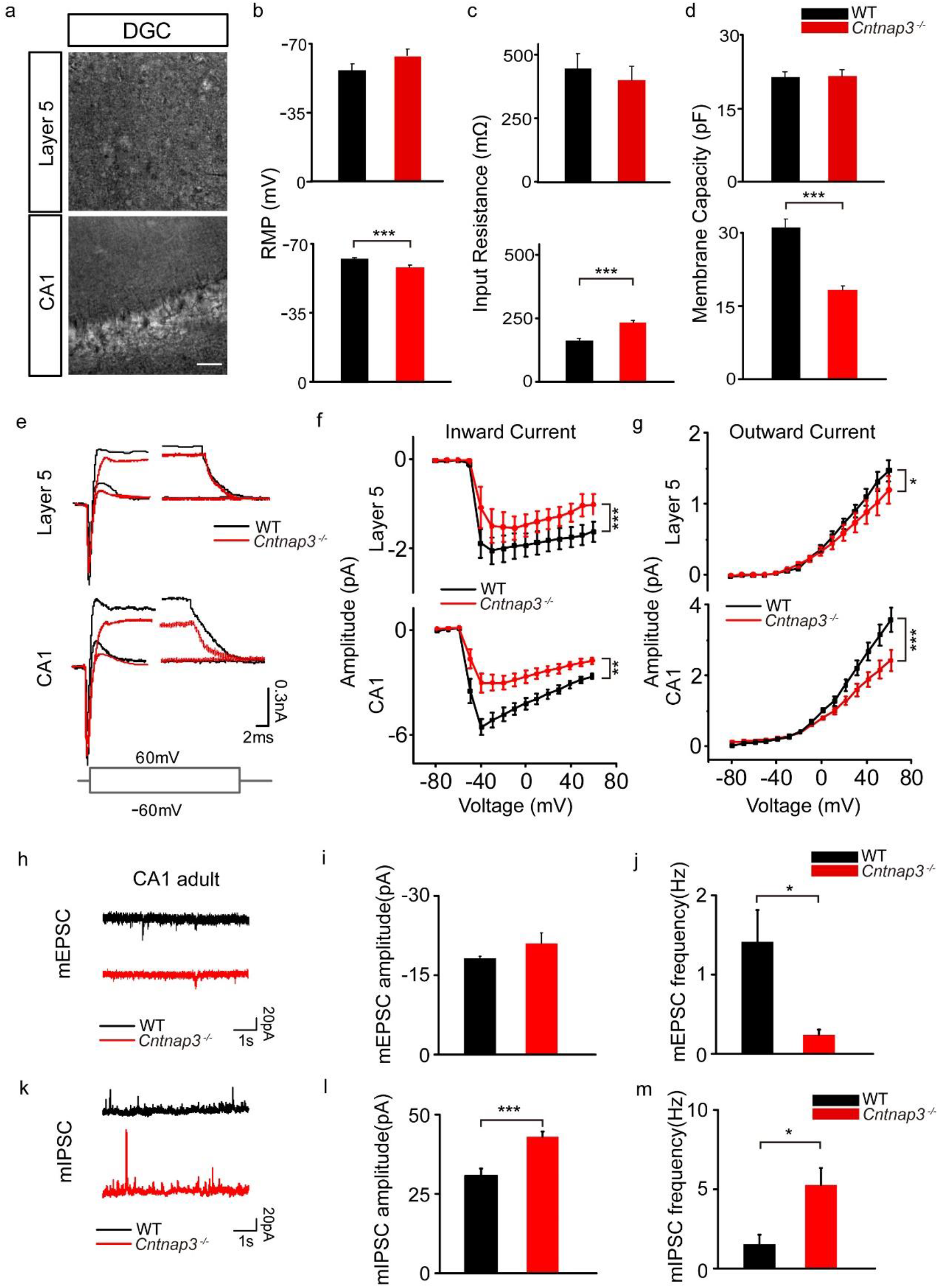
*Cntnap3^-/-^* mice exhibited decreased mEPSC and increased mIPSC frequency in hippocampal CA1 neurons. (a) Images of whole-cell recording of cortical layer 5 and hippocampal CA1 neurons in bright field, scale bars: 50 μm. Resting membrane potential (b), input resistance (c) and membrane capacity (d) of neurons in cortical layer 5 (upper panel, 29 cells from 3 WT mice, 26 cells from 3 KO mice) and hippocampal CA1 (lower panel, 38 cells from 4 WT mice, 39 cells from 4 WT mice). (e) Representative current responses of neurons in cortical layer 5 and hippocampal CA1 at -60mV and 60 mV. Amplitude of inward current (f) and outward current (g) responses of neurons in cortical layer 5 (upper panel, inward current: 26 traces from 3 WT mice, 21 traces from 3 KO mice, outward current: 29 traces from 3 WT mice, 26 traces from 3 KO mice) and hippocampal CA1 (lower panel, inward current: 32 traces from 4 WT mice, 33 traces from 4 KO mice, outward current: 38 traces from 4 mice, 33 traces from 4 KO mice) evoked by a series of voltage steps. (h) Representative mEPSC traces of neurons of hippocampal CA1 in *Cntnap3^-/-^* and WT adult mice. Amplitude (i) and frequency (j) of mEPSC of hippocampal CA1 in KO and WT adult mice (16 traces from 4 WT mice, 18 traces from 4 KO mice). (k) Representative mIPSC traces of neurons of hippocampal CA1 in KO and WT adult mice. Amplitude (l) and frequency (m) of mIPSC of hippocampal CA1 in KO and WT adult mice (20 traces from 4 WT mice, 20 traces from 4 KO mice). Statistical significance was evaluated by Student’s t test or two-way ANOVA (curve): * *p* <0.05; ** *p* <0.01; *** *p* <0.001. Error bars, ± SEM.

These developmental changes may further shape properties of action potentials and membrane activities. We thus investigated the inward and outward current during the excitation of neurons evoked by a series of voltage pulses in cortical layer 5 and hippocampal CA1 neurons of *Cntnap3^-/-^* and WT mice (Fig. 4e). We found that amplitude of inward currents of both layer 5 and CA1 neurons significantly decreased in *Cntnap3^-/-^* mice compared to those in WT mice (Fig. 4f). The outward currents displayed the same tendency (Fig. 4g). These results suggested that knockout of *Cntnap3* altered excitability of cortical and hippocampal neurons. However, when we measured firing rate and amplitude of action potentials of cortical layer 5 and hippocampal CA1 neurons, we found that these properties were not influenced by knockout of *Cntnap3* (Supplementary Fig. 7a-f). We reasoned that the altered excitability did not lead to change of fire rate may be due to the input of these neurons were also changed in the absence of *Cntnap3.*

Therefore, to elucidate the role of CNTNAP3 in synapse development, we further examine excitatory and inhibitory synaptic transmission in the hippocampal CA1 region of *Cntnap3^-/-^* mice. We measured miniature excitatory postsynaptic currents (mEPSC) and miniature inhibitory postsynaptic current (mIPSC) in the presence of tetrodotoxin (TTX) in acute hippocampal slices of *Cntnap3^-/-^* and WT mice (Supplementary Fig. 7g, h, i). We found that the frequency, but not amplitude, of mEPSC significantly decreased in adult *Cntnap3^-/-^* mice, in consistent with the observation of decreased excitatory synapse development *in vitro* and reduced spine density *in vivo* (Fig. 4i, j). Interestingly, both amplitude and frequency of mIPSC significantly increased in the hippocampal CA1 region of adult *Cntnap3^-/-^* mice, suggesting that numbers and strength of inhibitory synapses are both increased in *Cntnap3^-/-^* mice (Fig. 4l, m). We also measured mEPSC and mIPSC in adolescent mice from postnatal day 14 (P14) to 28 (P28) (Supplementary Fig. 7h, i). The frequency of mEPSC in *Cntnap3^-/-^* adolescent mice was significantly decreased (Supplementary Fig. 7j, k), whereas the frequency of mIPSC was markedly increased (Supplementary Fig. 7l, m). These results suggest that CNTNAP3 also play a pivotal role in regulating excitatory and inhibitory synaptic transmission *in vivo.* Taken together, these evidences indicate that CNTNAP3 regulates intrinsic neuronal excitability, as well as playing an opposite role in controlling development of excitatory and inhibitory synapses *in vivo.*

### Defects in social interaction, repetitive behaviors and cognition in *Cntnap3^-/-^* mice

Finally, we sought to determine whether deletion of *Cntnap3* may lead to autistic-like behavioral abnormalities in mouse, although previous studies showed that *Cntnap3* knockout mice exhibited the normal motor function and anxiety level(Hirata et al., 2016). We performed a battery of behavioral tasks on *Cntnap3^-/-^*, excitatory (*Nex-Cre: Cntnap3^flox/flox^*) and inhibitory (*Vgat-ires-Cre: Cntnap3^flox/flox^*) specific knockout mice. Remarkably, in the three-chamber test, *Cntnap3^-/-^* mice appeared no preference in staying with mouse over an empty cage, while still exhibited increased interaction time with novel mice over familiar mice, suggesting that *Cntnap3^-/-^* showed defects in social interactions (Fig. 5a, b, Supplementary Fig. 8a-c). Interestingly, we found that *Nex-Cre: Cntnap3^flox/flox^* mice, specific deletion of *Cntnap3* in excitatory neurons, showed no defects in recognizing either mice over cage, or novel over familiar mice (Fig. 5c, d, Supplementary Fig. 9a). However, inhibitory neuron specific knockout mice, *Vgat-ires-Cre: Cntnap3^flox/flox^*, exhibited defects in distinguishing novel mice with familiar mice, suggesting that *Cntnap3* in inhibitory neurons may contribute to social memory (Fig. 5e, f, Supplementary Fig. 9b).

**Figure 5.**
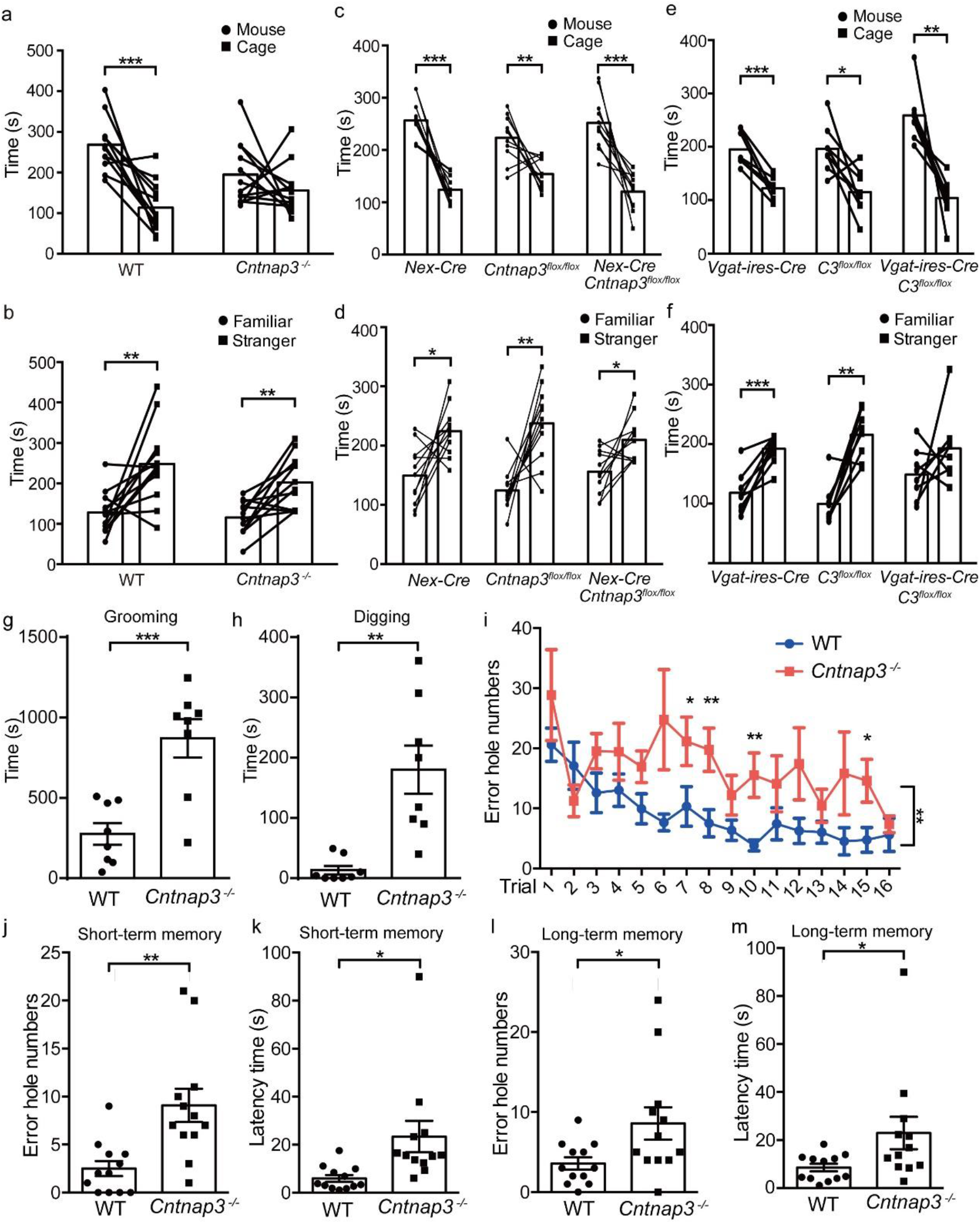
*Cntnap3* KO and conditional KO mice show abnormalities in social behavior, repetitive behaviors and learning memory tasks. Social behaviors in the three-chamber test (a-f), repetitive test (g,h), learning and memory task in the Barns maze task (i-m). (a) Time interacting with either an unfamiliar mouse or an empty cage within 10 min. (b) Time interacting with either a stranger mouse or a familiar mouse within 10 min. (n= 12 for either *Cntnap3*^-/-^ or WT mice). (c) Time interacting with either an unfamiliar mouse or an empty cage within 10 min. (d) Time interacting with either a stranger or a familiar mouse in 10 min. (n= 11 for each genotypes). (e) Time interacting with either an unfamiliar mouse or an empty cage within 10 min. (f) Time interacting with either a stranger mouse or a familiar mouse within 10 min. (n= 8 for each genotypes) (g) Time the mice spent to self-grooming in one hour within home cage. (h) Time the mice spent to digging in one hour within home cage. (n= 8 for either *Cntnap3^-/-^* or WT mice) (i) Learning curve as indicated by the error hole numbers before entering the hole during a 4-day training period (n= 12 for either *Cntnap3^-/-^* or WT mice). The error hole numbers (j) and latency time (up to 90s) (k) before entering the hole at the first test (short-term memory). The error hole numbers (l) and latency time (up to 90s) (m) before entering the hole at the second test (long-term memory). Statistical significance was evaluated by Student’s t test, paired *t* test (3 chamber tests) or two-way ANOVA (curve): *p < 0.05, ** *p* < 0.01, *** *p* < 0.001, Error bars, ± SEM.

Another phenotype of ASD patient is repetitive behaviors. Thus we measured the time which WT or *Cntnap3^-/-^* mice spent to groom or dig in one hour within its home cage. We found that *Cntnap3^-/-^* mice spent more time to self-groom or dig, which are considered to be repetitive behaviors (Fig. 5g, h, Supplementary Movie 1 and 2). These results suggest that *Cntnap3^-/-^* mice show the ASD-like behavior.

As the deletion of CNTNAP3 was found in the MR patient, we think that deletion of CNTNAP3 may lead to learning and memory disability, which is also an important accompanying phenotype of ASD. Next, we used the Barnes Maze test to examine whether *Cntnap3^-/-^* mice may have defects in learning and memory tasks. First, we found that *Cntnap3^-/-^* mice spent significantly longer time and more error holes to find the correct hole during the training session, suggesting that the learning ability in *Cntnap3^-/-^* mice was compromised (Fig. 5i, Supplementary Fig. 8d). We then performed the short-term and long-term memory tasks at day 5 and day 12 after training, respectively. We found that *Cntnap3^-/-^* mice spent markedly increased error holes and latency time in both memory tests, indicating the both short- and long-term memory capacity of *Cntnap3^-/-^* mice are severely affected (Fig. 5j-m, Supplementary Fig. 8e). We found no defects of *Cntnap3^-/-^* mice in consequent series of tests, including elevated plus maze (Supplementary Fig. 8f, g), light/dark shuttling (Supplementary Fig. 8h), open field test (Supplementary Fig. 8i, j), novel object recognition (Supplementary Fig. 8k), as well as context and cue-dependent fear conditioning (Supplementary Fig, 8l, m, n). These data suggest that *Cntnap3^-/-^* mice have specific defects related to social behaviors, repetitive behaviors, as well as learning and memory, but appeared normal levels of anxiety and fear conditioning responses.

In the excitatory neuron specific knockout mice (*Nex-Cre: Cntnap3^flox/flox^* and inhibitory neuron specific knockout mice (*Vgat-ires-Cre: Cntnap3^flox/flox^*), we found similar defects in learning curve of the Barnes Maze test (Fig. 6a-d), suggesting that proper functions of CNTNAP3 in both excitatory and inhibitory neurons are required for acquiring a certain type of spatial memory. Interestingly, deletion of CNTNAP3 in excitatory neurons exhibited more severe defects in long-term and short-term memory tasks, comparing to inhibitory neurons specific deletion (Fig. 6e-h, Supplementary Fig. 10a-d), suggesting that CNTNAP3 may play a more important role in glutamatergic neurons in regulating long-term memory maintenance. Similarly, we did not find any defects in the elevated plus maze and open field test in either excitatory neuron knockout (Supplementary Fig. 9c-f), or inhibitory neurons knockout mice (Supplementary Fig. 10e-h), suggesting that CNTNAP3 has no effects in regulating anxiety levels.

**Figure 6.**
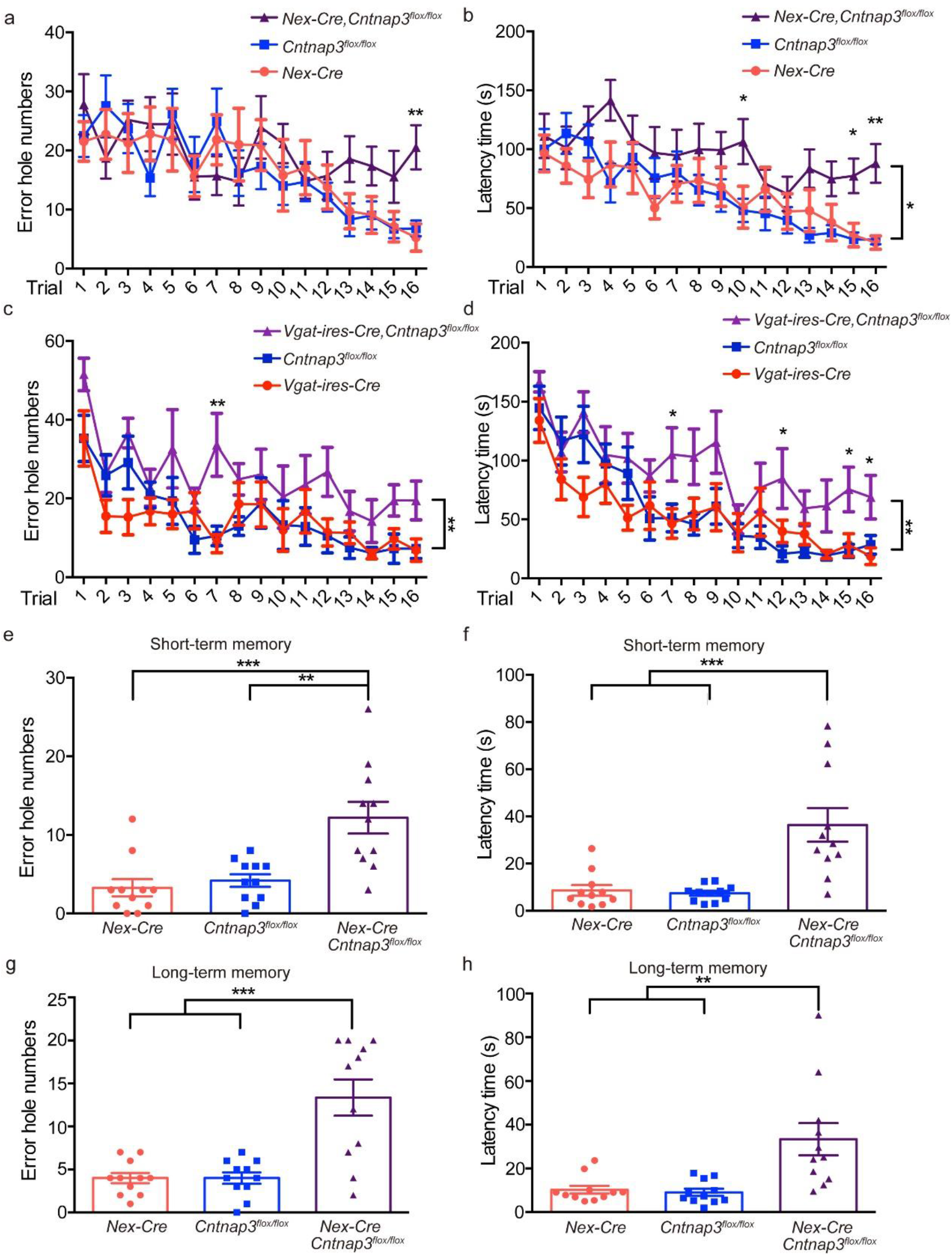
Both excitatory and inhibitory neuron specific deletion of *Cntnap3* exhibited abnormalities in learning and memory tasks. Learning curve as indicated by the error hole numbers to enter the hole during a 4-day training period (a, c) or indicated by the latency time (up to 180 s) before entering the hole during a 4-day training period (b, d). (a, b) *Nex-Cre, Cntnap3*^flox/flox^, (c,d) *vGat-ires-Cre, Cntnap3*^flox/flox^. (e-h) Short- and Long-term memory tasks for *Nex-Cre, Cntnap3*^flox/flox^ mice. (e) The error hole number before entering the hole at the first test (short-term memory). (f) The latency time before entering the hole (up to 90s) at the first test. (g) The error hole number before entering the hole at the second test (long-term memory). (h) The latency time before entering the hole (up to 90s) at the second test. Statistical significance was evaluated by one-way ANOVA, paired t test (3 chamber) or two-way ANOVA (curve): *p < 0.05, ** *p* < 0.01, *** *p* < 0.001, error bars, ± SEM.

## Discussion

We characterized critical functions of the synaptic adhesion molecule CNTNAP3 in synaptic development and transmission, as well as social behavior, repetitive behavior and cognitive tasks. The genetic connections between CNTNAP family and ASD are very intriguing over the years, as other synaptic adhesion molecules are implicated in ASD. R1219X was a stop-gain *de novo* mutations that was found in ASD patients, which indicated that CNTNAP3 might be a ASD candidate gene. In this study, we proved that R1219X could not interact with PSD-95 or Gephyrin, which were important scaffolding proteins in both excitatory synapses and inhibitory synapses. Moreover, R1219X could not rescue the neuronal morphology and synaptic changes which caused by knocking down *Cntnap3.* These suggest that R1219X may be a *loss-of-function* mutation of CNTNAP3 due to losing the function of its trans member region and intracellular part. Additionally, we identified two transmitted mutations (P614A, R786C) in *CNTNAP3* gene through whole-exome sequencing. Although neither mutation is a *de novo* mutation, we further addressed the functions of each mutations in regarding to synapse development and showed that they were indeed *loss-of-function* variants. But on the other hand, P614A and R786C are point mutations which only cause one amino acid mutation of CNTNAP3. Compairing with the stop-gain mutation of R1219X, the functional changes of P614A and R786C may be more lightly (Fig. 2g). That might be one reason why the fathers who also have the mutation of P614A or R786C are not ASD patients. Our results could only reveal the function of these mutations *in vitro*. As the in vivo system is more complicated, the lightly changes must be accumulated or induced by some other factors, for instance, the environmental factors, and then caused the ASD. These data suggested that besides *de novo* mutations, transmitted variants in ASD patients could also significantly contribute to pathophysiology of ASD.

Although *CNTNAP2* and *CNTNAP4* are ASD candidate genes in previous genetic and functional studies, the detail function of CNTNAPs in synapse development is yet to be determined. The finding that CNTNAP3 has interactions with NLGN1/2 family members strongly suggests that CNTNAP3 is an important synaptic adhesion molecule that plays critical roles in regulating synapse development, as confirmed by our experiments (Chih et al., 2005; Varoqueaux et al., 2006). Our data suggest that CNTNAP3 is an essential component of both excitatory synapse and inhibitory synapse. Further work is needed to determine the molecular mechanisms underlying the differentially regulatory function of CNTNAP3 in both excitatory and inhibitory synapses. The finding that CNTNAP3 regulates synapses development differentially provides novel insights for further understanding of the synapse basis for ASD.

Comparing with behavioral defects identified in *Cntnap2^-/-^* and *Cntnap4^-/-^* mice, we found rather specific defects in social behavior, repetitive behavior and cognitive tasks in *Cntnap3^-/-^* mice, suggesting that CNTNAP3 may play a more specific role in regulating social interaction, repetitive behavior, as well as learning and memory in mice. Further work of determining specific neural circuits responsible for defects in social behavior and learning memory would be crucial to provide insights into neural circuits regulating different cognitive behaviors specifically.

Combined the genetic mutations we discovered in Chinese Han ASD and Simons simplex collections, as well as functional evidences in genetic engineered mice, we suggest that *CNTNAP3* is an ASD candidate gene and the imbalance of excitatory and inhibitory synapse development caused by mutations of *CNTNAP3* contribute to the abnormal social behavior, repetitive behavior and cognitive functions.

## Materials and Methods

### Ethics statement

We obtained assent from the Institutional Review Board (IRB), Shanghai Mental Health Center of Shanghai Jiao Tong University (FWA Number: 00003065; IROG Number: 0002202). Dr. Yi-Feng Xu approved and signed our study with ethical review number 2016-4. Written informed consent was obtained from parents in consideration of the fact that all patients were minors. All participants were screened using the appropriate protocol approved by the IRB. Subjects’ information and Clinical Scale Assessment was described in previous report (22). The criteria of genetic testing are approved by the ethics committees of Children’s Hospital, Fudan University (2014-107). Pre-test counseling were performed by physicians, appropriate informed consents were signed by patient’s parents in clinics.

### Simons simplex collection data

We are grateful to all of the families at the participating Simons Simplex Collection (SSC) sites, as well as the principal investigators (A. Beaudet, R. Bernier, J. Constantino, E. Cook, E. Fombonne, D. Geschwind, R. Goin-Kochel, E. Hanson, D. Grice, A. Klin, D. Ledbetter, C. Lord, C. Martin, D. Martin, R. Maxim, J. Miles, O. Ousley, K. Pelphrey, B. Peterson, J. Piggot, C. Saulnier, M. State, W. Stone, J. Sutcliffe, C. Walsh, Z. Warren, E. Wijsman).

Approved researchers can obtain the SSC population dataset described in this study (https://simons.wuxinextcode.com/csa/welcome) by applying at https://base.sfari.org.

### In situ hybridization

Cut 25 g of CNTNAP3 ISH plasmid DNA (T Vector, Promega, A3600) at the 5’ end with Sal1 enzyme (NEB, R3138G) for 8 hours. Then transcript of mRNA probe with T7 RNA polymerase (Promega, P2077). Sections (30 m) were postfixed in 4% DEPC-PFA at 4oC overnight. After wash, the sections were acetylated for 10 mins on the rocker with Acetylation mix. Pre-hybridize for 2 hours in hybridization solution in a 65 oC water bath before the 16 hours hybridization with 1 μg/ml of probe. After wash the sections were incubated overnight at 4 oC in 1:2000 anti-digoxigenin antibody (Roche, 11093274910). After wash, stain the sections in fast red solution (HNPP fluorescent detection set, Roche, 11758888001) for more than 3 hours. Images were obtained using a confocal microscope (Nikon TiE, Olympus FV10i).

### RNA isolation and RTq-PCR

Tissues of mice were separated and collected in Trizol (Invitrogen, 15596-018). cDNAs were obtained by Reverse Transcriptase M-MLV kit (TaKaRa, D2639B). mRNA levels of CNTNAP3 were examined by SYBR green (Toyobo, QPK-201)-based quantitative real time PCR (qPCR). qPCR was performed on a ABI-Gene sequence detection system. The GAPDH gene was used as control. Forward and reverse primers used were as following: m-CNTNAP3: forward: TCAAAACCGGGCTAGAGTCA reverse: GGGGCTGTCTATCTTCTGCG m-GAPDH: forward: ACGGCCGCATCTTCTTGTGCAGTG reverse: GGCCTTGACTGTGCCGTTGA

### 293T cell culture and co-immunoprecipitation

293T cell was cultured in the DMEM (Gibco, 11965-092) medium with 10% FBS. The cells were transfected with Lipo-2000 (Invitrogen, 11680-019). After 2 days of transfection, the cells were lysed in RIPA buffer (1%Triton, 150 mM NaCl, 25 mM Tris-Hcl (PH=7.4), 2 mM EDTA (PH=8.0)) for 1.5h at 4°C. After the Sonic broken of 10 min and 5000 rpm centrifuge for 10 min, the supernatant was added with protein A/G agarose (Abmart, A10001M) with IgG or anti-GFP (Abmart, A20004L), anti-HA-tag mouse mAb agarose conjugated (Abmart, M20013S) or anti-myc-tag mouse mAb agarose conjugated (Abmart, M20012S) for 4 hours. Wash the agarose conjugated beads 3 times before loading the protein loading.

### Western blot

8% acrylamide was used in the western blot. The 1st antibody was incubated overnight at 4 °C and the 2nd antibody was incubated for 2 hours. Antibodies used were as following:

Anti-HA: 1:1000 Covance, MMS-101R

Anti-Myc: 1:1000 Abmart, M2002L

Anti-GFP: 1:1000 Abmart, A20004L

Sheep anti mouse HRP, GE-Healthcare, NA931V

### Neuron culture and calcium phosphate transfection

Cultures were prepared from embryonic day 13.5 WT C57BL/6 mice. The cells form cortex were separated with papain (Worthington, LS003126) and then cultured in the Neurobasal medium (Gibco, 21103-049) with 0.2% B27 (Gibco, 17504-044).

The neurons were transfected on the day 5 by calcium phosphate transfection. 2μg of each plasmids was used to do the transfection. At the day 14, the neurons were collected to do the immunohistochemical staining.

### Genetic engineered mice

Cntnap3 knockout and Cntnap3 conditional knockout mice were customized designed and ordered from Biocytogen.

### Immunohistochemical staining

For the cultured neurons, the neurons were fixed in 4% PFA at room temperature for 20 min before wash. The 1st antibody was incubated overnight at 4°C and the 2nd antibody was incubated at room temperature for 2 hours. For the brain slice, the slice was cut in 40μm. The 1st antibody was incubated overnight at 4°C and the 2nd antibody was incubated at room temperature for 2 hours. Antibodies used were as following:

GFP (Rabbit): 1:1000 Invitrogen, A11122

GFP (Mouse): 1:1000 Earthox, E022030-10

vGlut1: 1:3000 Millipore, AB5905

vGat: 1:1000 Synaptic Systerms, 31002

PV: 1:1000 Millipore, MAB157

SST: 1:250 Santa Cruz, SC-7819

Donkey anti mouse CF488A conjugate, Biotium, 20014

Donkey anti rabbit CF488A conjugate, Biotium, 20015

Donkey anti mouse CF555A conjugate, Biotium, 20037

Goat anti guinea pig CF555A conjugate, Biotium, 20036

Donkey anti rabbit CF647A conjugate, Biotium, 20047

Images were obtained using a confocal microscope (Nikon TiE, Nikon A1R, Nikon NiE, Olympus FV10i)

### Golgi Staining

The FD Rapid GolgiStain Kit (FD neuro technologies, PK-401) was used for the Golgi staining. The mouse brains were put in the A&B mixed medium for 14 days and then in the solution C for 5 days. The slice was cut with 100μm. Images were obtained using a confocal microscope (Olympus FV1000).

### Behavior test

All the mice used in this study were male and handled for more than 3 days prior to behavioral tasks. The animals’ movements were recorded and analysed by Ethovision XT software (Noldus, Wageningen, Netherlands). The equipment used in the behavior test was thoroughly cleaned with 75% isopropanol before each mouse was tested.

### Open field

The behavior was recorded for 30 min. The mouse was put in the 45cm×45cm box for free moving.

### Elevated plus maze

The plus maze with 2 close arms and 2 open arms was used in the experiment. The mouse was placed in the center of the maze with its head to the close arm to start the experiment and then recorded for 10 min.

### Three chamber test

The box with 3 chambers was used in the test. The test mouse was put in the center chamber for 3 times test (10 min each). For the first time, both sides of the chambers were empty. For the second time, one WT mouse was put in one random cage, the other cage was empty. For the last time, one stranger WT mouse was added to the leftover empty cage. In all of the 3 times of test, the test mouse was free moving. Only the mouse in the interaction zone (about 8cm beside the cage) was used as the interaction time.

### Barnes maze

A disc with 40 holes was used in the experiment. At the learning period, animals will receive 4 trials per day with an inter-trial interval of 15 minutes during 4 days. The short-term memory test was done on the day 5 and the long-term memory test done was on the day 12.

During the learning test, the time is 3 min, if the mouse failed to enter the escape cage; we helped it enter the cage and recorded the latency time with 180s. At the memory test, the time is 90s without escape cage. During all of the test, we use glare as the stimulation.

### The light and dark test

The box with 2 chambers was used in the test. One of the chamber was covered with a black plate which made the box inside dark, the other chamber was empty. The box was put under the light and the mouse was put into the dark part when the test started. The time mouse spend in the light area was recorded in the 10 minutes test.

### Novel object recognition

At the first test, two toys with the same size were put into the opposite sides of the open field box. The mouse was put in the center of the box for free moving in 10 minutes. At the second test, one of the toy was changed by a stranger toy. The time mouse spend with the stranger toy was recorded in the 10 minutes test.

### Fear conditioning test

0.8mA, 3 tries were used in the fear conditioning test. The freeze percentage was test in both contextual and cue fear. The voice was used only without shock was used in the extinction test.

### Home cage video

The mouse was put in the home cage 2 days before the video started. The video was recorded for 24hours. The first half hour after turn off the light and the half hour 6 hours after turn off the light were counted. All of the experiments were in the normal circadian rhythms (12h light and 12h dark).

### The mouse CNTNAP3 RNAi sequence

CAGACAGTGTGGTACAATA

### Electrophysiological Recordings in Slices

Mice were anaesthetized with avertin and decapitated. The brain was quickly removed and submerged in oxygenated (95% O2 and 5% CO2) ice-cold sucrose-based artificial cerebrospinal fluid (sucrose-based aCSF) containing (in mM: 234 Sucrose, 2.5 KCl, 26 NaHCO3, 1.25 NaH2PO4, 11 D-Glucose, 0.5 CaCl2 and 10 MgSO4). Coronal slices (300 μm) were prepared using a vibratome (VT1200S, Leica) and kept in an incubating chamber filled with oxygenated aCSF (in mM: 126 NaCl, 3 KCl, 26 NaHCO3, 1.2 NaH2PO4, 10 D-Glucose, 2.4 CaCl2 and 1.3 MgCl2) at 34°C. After a recovery period at least 40 minutes, an individual slice was transferred to a recording chamber and was continuously superfused with oxygenated aCSF at a rate of 3-5 ml per minute at 30 ± 1 °C. An inverted microscope (Andor) equipped with epifluorescence and infrared-differential interference contrast (DIC) illumination, a camera, and one air immersion lense (4X) and one water immersion lense (40X) were used to visualize and target recording electrodes and cells.

Whole-cell patch clamp recording was performed on cells from the layer 5 of neocortex and CA1 of hippocampus. Patch pipettes had a 5-7 MΩ resistance when filled with intracellular solution (in mM: 130 potassium gluconate, 16 KCl, 2 MgCl2, 10 HEPES, 0.2 EGTA, 4 Na2-ATP, 0.4 Na3-GTP, pH = 7.25, adjusted with KOH). Miniature EPSCs and IPSCs were recorded by holding the membrane potential at -70 mV and 12 mV after the treatment of 2.5 mM TTX. Inward currents were recorded in voltage-clamp mode with a basal holding potential of -60 mV followed by stimulating pulses from -80 mV to 60 mV with a step size of 10 mV. Evoked action potentials were recorded in current-clamp mode using a series of depolarizing currents ranged from -100 pA to 280 pA in increments of 20 pA.

Evoked EPSCs and IPSCs were recorded using record glass pipettes as above (5 to 7 MΩ) and were evoked using a concentric bipolar electrode (WPI; FHC, CBBEB75) which was used to stimulate neurons with a brief current pulse (50 ms), and the current pulse was delivered by a stimulus isolation unit (ISO-Flex, A.M.P.I., Jerusalem, Israel). Signals were filtered at 2 kHz and digitized at 100 kHz using Digidata 1440A (Molecular Devices). Data acquisition and slope measurement were carried out using pClamp 10.2 (Molecular Devices). Pulse generation was achieved using a Master 8 stimulator (A.M.P.I., Jerusalem, Israel).

## Acknowledgements

We thank the team of WuXi NextCODE for technical assistance in SSC data analysis. This work was supported by the National Basic Research Program of China (2014CB964600, 2017YFA0105201, 2017YFA0103303, 2017YFA0102601), NSFC Grants (#31625013, #91732302, #81527901, #81471484, #31670842). the Strategic Priority Research Program of the Chinese Academy of Sciences (XDA01020309), and Shanghai Brain-Intelligence Project from STCSM (16JC1420500). The research is supported by the Open Large Infrastructure Research of Chinese Academy of Sciences. We appreciate obtaining access to genotype, phenotype and pedigree data on SFARI Base.

## AUTHOR CONTRIBUTIONS

Z.Q designed the study. D.T performed most of the experiment and data analysis. R.C, L.H, X.W performed the electrophysiology experiment. J.L, X.X contributed to *in situ* hybridation experiments. Y.L, W.Z. contributed to the sample collection and NGS analysis for R786C mutation. Y.Z contributed in the mice feeding and genotyping. J S.S contributed in the cell culture. Y.Z. and Y.S.D contributed the sample collection and NGS analysis for P614A mutations. D.T, R.C and Z.Q wrote the manuscript with the help of all other authors.

## References

Alarcon, M., Abrahams, B.S., Stone, J.L., Duvall, J.A., Perederiy, J.V., Bomar, J.M., Sebat, J., Wigler, M., Martin, C.L., and Ledbetter, D.H., et al. (2008). Linkage, association, and gene-expression analyses identify CNTNAP2 as an autism-susceptibility gene. AM J HUM GENET 82, 150–159.

Anderson, G.R., Galfin, T., Xu, W., Aoto, J., Malenka, R.C., and Sudhof, T.C. (2012). Candidate autism gene screen identifies critical role for cell-adhesion molecule CASPR2 in dendritic arborization and spine development. Proc Natl Acad Sci U S A 109, 18120–18125.

Arking, D.E., Cutler, D.J., Brune, C.W., Teslovich, T.M., West, K., Ikeda, M., Rea, A., Guy, M., Lin, S., and Cook, E.H., et al. (2008). A common genetic variant in the neurexin superfamily member CNTNAP2 increases familial risk of autism. AM J HUM GENET 82, 160–164.

Ashrafi, S., Betley, J.N., Comer, J.D., Brenner-Morton, S., Bar, V., Shimoda, Y., Watanabe, K., Peles, E., Jessell, T.M., and Kaltschmidt, J.A. (2014). Neuronal Ig/Caspr recognition promotes the formation of axoaxonic synapses in mouse spinal cord. NEURON 81, 120–129.

Bakkaloglu, B., O’Roak, B.J., Louvi, A., Gupta, A.R., Abelson, J.F., Morgan, T.M., Chawarska, K., Klin, A., Ercan-Sencicek, A.G., and Stillman, A.A., et al. (2008). Molecular cytogenetic analysis and resequencing of contactin associated protein-like 2 in autism spectrum disorders. AM J HUM GENET 82, 165–173.

Bellen, H.J., Lu, Y., Beckstead, R., and Bhat, M.A. (1998). Neurexin IV, caspr and paranodin--novel members of the neurexin family: encounters of axons and glia. TRENDS NEUROSCI 21, 444–449.

Chih, B., Engelman, H., and Scheiffele, P. (2005). Control of excitatory and inhibitory synapse formation by neuroligins. SCIENCE 307, 1324–1328.

de la Torre-Ubieta, L., Won, H., Stein, J.L., and Geschwind, D.H. (2016). Advancing the understanding of autism disease mechanisms through genetics. NAT MED 22, 345–361.

Feng, J., Schroer, R., Yan, J., Song, W., Yang, C., Bockholt, A., Cook, E.J., Skinner, C., Schwartz, C.E., and Sommer, S.S. (2006). High frequency of neurexin 1beta signal peptide structural variants in patients with autism. NEUROSCI LETT 409, 10–13.

Hirata, H., Takahashi, A., Shimoda, Y., and Koide, T. (2016). Caspr3-Deficient Mice Exhibit Low Motor Learning during the Early Phase of the Accelerated Rotarod Task. PLOS ONE 11, e147887.

Huguet, G., Ey, E., and Bourgeron, T. (2013). The genetic landscapes of autism spectrum disorders. Annu Rev Genomics Hum Genet 14, 191–213.

Jamain, S., Quach, H., Betancur, C., Rastam, M., Colineaux, C., Gillberg, I.C., Soderstrom, H., Giros, B., Leboyer, M., and Gillberg, C., et al. (2003). Mutations of the X-linked genes encoding neuroligins NLGN3 and NLGN4 are associated with autism. NAT GENET 34, 27–29.

Karayannis, T., Au, E., Patel, J.C., Kruglikov, I., Markx, S., Delorme, R., Heron, D., Salomon, D., Glessner, J., and Restituito, S., et al. (2014). Cntnap4 differentially contributes to GABAergic and dopaminergic synaptic transmission. NATURE 511, 236–240.

Kashani, A.H., Qiu, Z., Jurata, L., Lee, S.K., Pfaff, S., Goebbels, S., Nave, K.A., and Ghosh, A. (2006). Calcium activation of the LMO4 transcription complex and its role in the patterning of thalamocortical connections. J NEUROSCI 26, 8398–8408.

Kim, H.G., Kishikawa, S., Higgins, A.W., Seong, I.S., Donovan, D.J., Shen, Y., Lally, E., Weiss, L.A., Najm, J., and Kutsche, K., et al. (2008). Disruption of neurexin 1 associated with autism spectrum disorder. AM J HUM GENET 82, 199–207.

Kong, S.W., Collins, C.D., Shimizu-Motohashi, Y., Holm, I.A., Campbell, M.G., Lee, I.H., Brewster, S.J., Hanson, E., Harris, H.K., and Lowe, K.R., et al. (2012). Characteristics and predictive value of blood transcriptome signature in males with autism spectrum disorders. PLOS ONE 7, e49475.

Mosrati, M.A., Schrauwen, I., Kamoun, H., Charfeddine, I., Fransen, E., Ghorbel, A., Van Camp, G., and Masmoudi, S. (2012). Genome wide analysis in a family with sensorineural hearing loss, autism and mental retardation. GENE 510, 102–106.

Peles, E., Nativ, M., Lustig, M., Grumet, M., Schilling, J., Martinez, R., Plowman, G.D., and Schlessinger, J. (1997). Identification of a novel contactin-associated transmembrane receptor with multiple domains implicated in protein-protein interactions. EMBO J 16, 978–988.

Penagarikano, O., Abrahams, B.S., Herman, E.I., Winden, K.D., Gdalyahu, A., Dong, H., Sonnenblick, L.I., Gruver, R., Almajano, J., and Bragin, A., et al. (2011). Absence of CNTNAP2 leads to epilepsy, neuronal migration abnormalities, and core autism-related deficits. CELL 147, 235–246.

Poliak, S., Salomon, D., Elhanany, H., Sabanay, H., Kiernan, B., Pevny, L., Stewart, C.L., Xu, X., Chiu, S.Y., and Shrager, P., et al. (2003). Juxtaparanodal clustering of Shaker-like K+ channels in myelinated axons depends on Caspr2 and TAG-1. J CELL BIOL 162, 1149–1160.

Rios, J.C., Melendez-Vasquez, C.V., Einheber, S., Lustig, M., Grumet, M., Hemperly, J., Peles, E., and Salzer, J.L. (2000). Contactin-associated protein (Caspr) and contactin form a complex that is targeted to the paranodal junctions during myelination. J NEUROSCI 20, 8354–8364.

Spiegel, I., Salomon, D., Erne, B., Schaeren-Wiemers, N., and Peles, E. (2002). Caspr3 and caspr4, two novel members of the caspr family are expressed in the nervous system and interact with PDZ domains. MOL CELL NEUROSCI 20, 283–297.

Szatmari, P., Paterson, A.D., Zwaigenbaum, L., Roberts, W., Brian, J., Liu, X.Q., Vincent, J.B., Skaug, J.L., Thompson, A.P., and Senman, L., et al. (2007). Mapping autism risk loci using genetic linkage and chromosomal rearrangements. NAT GENET 39, 319–328.

Traka, M., Goutebroze, L., Denisenko, N., Bessa, M., Nifli, A., Havaki, S., Iwakura, Y., Fukamauchi, F., Watanabe, K., and Soliven, B., et al. (2003). Association of TAG-1 with Caspr2 is essential for the molecular organization of juxtaparanodal regions of myelinated fibers. J CELL BIOL 162, 1161–1172.

Traut, W., Weichenhan, D., Himmelbauer, H., and Winking, H. (2006). New members of the neurexin superfamily: multiple rodent homologues of the human CASPR5 gene. MAMM GENOME 17, 723–731.

Turner, T.N., Coe, B.P., Dickel, D.E., Hoekzema, K., Nelson, B.J., Zody, M.C., Kronenberg, Z.N., Hormozdiari, F., Raja, A., and Pennacchio, L.A., et al. (2017). Genomic Patterns of De Novo Mutation in Simplex Autism. CELL 171, 710–722.

Vaags, A.K., Lionel, A.C., Sato, D., Goodenberger, M., Stein, Q.P., Curran, S., Ogilvie, C., Ahn, J.W., Drmic, I., and Senman, L., et al. (2012). Rare deletions at the neurexin 3 locus in autism spectrum disorder. AM J HUM GENET 90, 133–141.

Varoqueaux, F., Aramuni, G., Rawson, R.L., Mohrmann, R., Missler, M., Gottmann, K., Zhang, W., Sudhof, T.C., and Brose, N. (2006). Neuroligins determine synapse maturation and function. NEURON 51, 741–754.

Wen, Z., Cheng, T.L., Li, G.Z., Sun, S.B., Yu, S.Y., Zhang, Y., Du YS, and Qiu, Z. (2017). Identification of autism-related MECP2 mutations by whole-exome sequencing and functional validation. MOL AUTISM 8, 43.

Willsey, A.J., and State, M.W. (2015). Autism spectrum disorders: from genes to neurobiology. CURR OPIN NEUROBIOL 30, 92–99.

